# Genome-wide association study suggests an independent genetic basis of zinc and cadmium concentrations in fresh sweet corn kernels

**DOI:** 10.1101/2021.02.19.432009

**Authors:** Matheus Baseggio, Matthew Murray, Di Wu, Gregory Ziegler, Nicholas Kaczmar, James Chamness, John P. Hamilton, C. Robin Buell, Olena K. Vatamaniuk, Edward S. Buckler, Margaret E. Smith, Ivan Baxter, William F. Tracy, Michael A. Gore

**Affiliations:** Plant Breeding and Genetics Section, School of Integrative Plant Science, Cornell University, Ithaca, NY 14853, USA; Department of Agronomy, University of Wisconsin-Madison, Madison, WI, 53706 USA; Donald Danforth Plant Science Center, St. Louis, MO 63132, USA; Department of Plant Biology, Michigan State University, East Lansing, MI 48824, USA; Soil and Crop Sciences Section, Plant Biology Section, School of Integrative Plant Science, Cornell University, Ithaca, NY 14853 USA; Institute for Genomic Diversity, Cornell University, Ithaca, NY 14853 USA; US Department of Agriculture-Agricultural Research Service, Robert W. Holley Center for Agriculture and Health, NY 14853 USA

**Keywords:** genome-wide association study, whole-genome prediction, elements, kernels, sweet corn

## Abstract

Despite being one of the most consumed vegetables in the United States, the elemental profile of sweet corn (*Zea mays* L.) is limited in its dietary contributions. To address this through genetic improvement, a genome-wide association study was conducted for the concentrations of 15 elements in fresh kernels of a sweet corn association panel. In concordance with mapping results from mature maize kernels, we detected a probable pleiotropic association of zinc and iron concentrations with *nicotianamine synthase5* (*nas5*), which purportedly encodes an enzyme involved in synthesis of the metal chelator nicotianamine. Additionally, a pervasive association signal was identified for cadmium concentration within a recombination suppressed region on chromosome 2. The likely causal gene underlying this signal was *heavy metal ATPase3* (*hma3*), whose counterpart in rice, *OsHMA3*, mediates vacuolar sequestration of cadmium and zinc in roots, whereby regulating zinc homeostasis and cadmium accumulation in grains. In our association panel, *hma3* associated with cadmium but not zinc accumulation in fresh kernels. This finding implies that selection for low cadmium will not affect zinc levels in fresh kernels. Although less resolved association signals were detected for boron, nickel, and calcium, all 15 elements were shown to have moderate predictive abilities via whole-genome prediction. Collectively, these results help enhance our genomics-assisted breeding efforts centered on improving the elemental profile of fresh sweet corn kernels.

## INTRODUCTION

As with all multicellular organisms, the concentration and distribution of elements in tissues and organs influence growth and development over the plant life cycle. At least 16 elements (boron, calcium, carbon, chlorine, copper, hydrogen, iron, magnesium, manganese, molybdenum, nitrogen, oxygen, phosphorus, potassium, sulfur, and zinc) are considered essential for higher plant species, with an additional four elements (cobalt, silicon, nickel, and sodium) essential for a subset of higher plants (Mengel and Kirkby 2001). In plants, the need for and concentration of macroelements (carbon, hydrogen, oxygen, nitrogen, phosphorus, sulfur, potassium, calcium, magnesium) (Hawkesford *et al*. 2012) are relatively greater than for microelements (iron, manganese, copper, zinc, nickel, molybdenum, boron, chlorine, and cobalt) (Broadley *et al*. 2012). Additionally, nonessential heavy metals such as cadmium, chromium, and lead that lack involvement in normal physiological functions can accumulate to toxic concentrations in plants, penetrating the food chain and posing a threat to human health (Singh *et al*. 2016).

Not unlike plants, humans can suffer a range of adverse health effects from the excess or deficiency of essential and nonessential elements. Of all the micronutrients, deficiency is most prevalent for iron, with more than two billion people affected worldwide (Viteri 1998). Of similar scale, nearly two billion people are estimated to suffer from dietary zinc deficiency throughout developing nations (Prasad 2014). Given that metal chelating substances such as phytate in cereal grains bind zinc and iron and inhibit their absorption, zinc and iron deficiencies resulting from low bioavailability could coincide (Sandstead and Smith 1996; Lönnerdal 2000). Although severe dietary micronutrient deficiency is far less prevalent in developed nations, approximately 10 million people are iron deficient in the U.S. (Miller 2013). In the U.S., the average daily intake of iron by most premenopausal (12 mg d^−1^) and pregnant (15 mg d^−1^) women is 6 and 12 mg d^−1^ less than its recommended daily allowance (RDA), respectively (Institute of Medicine 2001; Linus Pauling Institute 2016). Additionally, the zinc RDA for adult women and men is 11 mg d^−1^ and 8 mg d^−1^, with elderly in the U.S. at higher risk for mild zinc deficiency (Mocchegiani *et al*. 2013; Linus Pauling Institute 2019).

Crop biofortification via genomics-assisted breeding and genetic engineering has emerged as an attractive approach for nutritional enhancement of crops and the generation of new varieties with a high density of iron and zinc in edible plant tissues (Murgia *et al*. 2012; Bhullar and Gruissem 2013; Hirschi 2020). It is increasingly recognized that plant membrane transporters and metal chelators are among the key targets for increasing mineral nutrient density in plant tissues (Waters and Sankaran 2011; Schroeder *et al*. 2013). However, in many cases, transporters that facilitate the accumulation of iron and zinc are multispecific and can mediate the uptake and internal transport of nonessential and potentially highly toxic heavy metals including cadmium (Waters and Sankaran 2011; Schroeder *et al*. 2013; Khan *et al*. 2014). Therefore, efforts to increase the concentration of iron and zinc in grains of cereals could also increase the concentration of cadmium. This poses a serious threat to food security, especially if crops are grown on soils either contaminated with cadmium or low in microelements, particularly iron (Waters and Sankaran 2011; Schroeder *et al*. 2013; Khan *et al*. 2014).

Vegetative and seed tissues of fruits and vegetables are important dietary sources of essential and nonessential elements for humans to meet their daily nutrient needs. Given that sweet corn is the third most consumed vegetable in the U.S. (USDA-NASS 2018), the elemental profile of fresh sweet corn kernels is an important consideration for human health and nutrition. Although not a major contribution to the RDA of iron and zinc, the consumption of 100 g of uncooked, yellow sweet corn (medium-sized ear) provides 0.52 mg of iron and 0.46 mg of zinc (USDA-ARS 2019), but the bioavailable amount is expected to be no more than a quarter of the total of each element (Bouis and Welch 2010). Therefore, there exists a tremendous opportunity to improve the elemental profile of fresh sweet corn kernels through genomics-assisted breeding, but this first requires an understanding of the phenotypic variability and genetic control of zinc, iron, and other elements. Considerable heritable variation exists for elemental concentrations in physiologically mature grain of diverse maize panels (Ziegler *et al*. 2017; Wu *et al*. 2021), but a comparable level of genetic understanding is severely lacking for immature kernels (fresh-eating stage) of diverse sweet corn germplasm.

Complex physiological and genetic networks coordinate elemental uptake, transport, and accumulation in plants, and these processes are responsive to the environment in which plants are grown (Reviewed in Baxter 2009). In the genomes of Arabidopsis, maize, rice, and other plant species, gene families have been identified for metal transporters and chelators including but not limited to HEAVY-METAL ATPASE (HMA), OLIGOPEPTIDE TRANSPORTERS (OPTs) and their subfamily of YELLOW STRIPE-LIKE (YSL), ZINC-REGULATED TRANSPORTER (ZRT)/IRON-REGULATED TRANSPORTER (IRT)-LIKE PROTEIN (ZIP) and NICOTIANAMINE SYNTHASE (NAS) (Whitt *et al*. 2020). As it relates to maize, *yellow stripe1* (*ys1*) and *ys3* encode proteins that have been functionally shown to transport iron when associated with a strong metal-ligand, nicotianamine— synthesized from *S*-adenosyl-methionine by NAS enzymes—or its derivative phytosiderophores such as mugineic acid and deoxymugenic acid (Von Wiren *et al*. 1994; Chan-Rodriguez and Walker 2018), whereas the proteins encoded by *ysl2* and *zip5* have been functionally implicated in the accumulation of zinc and iron in grain (Li *et al*. 2019; Zang *et al*. 2020). Despite these advancements, the vast majority of metal transporters and chelators in the maize genome have not been deeply characterized at the functional level, thus a wide gap in the knowledge base remains for the key gene family members controlling the content and composition of elements in maize tissues and organs.

Explaining and predicting the quantitative variation of phenotypes is a major challenge in crop plants, but there has been notable recent progress for maize grain elemental phenotypes (Ziegler *et al*. 2017; Hindu *et al*. 2018; Wu *et al*. 2021). In the U.S. maize nested association mapping (NAM) panel, joint-linkage analysis and genome-wide association studies (GWAS) were used to identify six strong candidate genes for the concentrations of manganese, molybdenum, phosphorus, or rubidium in physiologically mature grain (Ziegler *et al*. 2017). Through the implementation of GWAS in the maize Ames panel, Wu *et al*. (2021) resolved several loci previously identified to control variation for copper, iron, manganese, molybdenum, and/or zinc in mature grain from the U.S. NAM panel, which resulted in the identification of two metal chelator and five metal transporter candidate genes. Additionally, the authors detected novel candidate gene loci for boron and nickel grain concentrations. Whole-genome prediction (WGP) models have been found to be moderately predictive of elemental concentrations in mature grain of tropical maize populations (zinc) (Guo *et al*. 2020; Mageto *et al*. 2020) and the Ames panel (boron, calcium, copper, iron, potassium, magnesium, manganese, molybdenum, nickel, phosphorus, and zinc) (Wu *et al*. 2021). Notwithstanding this progress with mature grain, the genotype-phenotype map of elemental concentrations in fresh sweet corn kernels is completely nonexistent, thus there exists tremendous opportunities for studying the quantitative genetics of these nutritionally relevant phenotypes.

In this study, we used a sweet corn association panel for the genetic dissection and prediction of quantitative variation of 15 elements in fresh sweet corn kernels. The three major objectives of our study were to (i) evaluate variability and heritability of elemental fresh kernel phenotypes within and across field locations, (ii) employ GWAS to identify candidate genes associated with the levels of elements in fresh kernels, and (iii) assess the predictive abilities of WGP models as an evaluation of the potential that genomic selection has for the genetic improvement of elemental fresh kernel phenotypes important to human nutrition and health.

## MATERIALS AND METHODS

### Plant Materials and Experimental Design

In two consecutive field seasons (2014-15), a sweet corn association panel of 430 inbred lines representing the genetic diversity of temperate U.S. breeding programs (Baseggio *et al*. 2019) was evaluated at Cornell University’s Musgrave Research Farm in Aurora, NY, on a Lima silt loam (fine-loamy, mixed, semiactive, mesic Oxyaquic Hapludalfs) and University of Wisconsin’s West Madison Research Station in Verona, WI, on a Plano silt loam (fine-silty, mixed, superactive, mesic Typic Argiudolls). The panel consists of *sugary1* (*su1*), *sugary1:sugary enhancer1* (*su1se1*), *shrunken2* (*sh2*), *sugary1:shrunken2* (*su1sh2*), *brittle2* (*bt2*), and *amylose-extender:dull:waxy* (*aeduwx*) lines that are homozygous for endosperm mutations that cause deficiences in starch biosynthesis. Additionally, there were 20 non-sweet corn inbred lines and four repeated check sweet corn inbred lines included in the experiment. In each of the four environments (location × year combination), the experiment was arranged as an augmented incomplete block design grown as a single replicate as previously described by Baseggio *et al*. (2019). Briefly, the lines were separated into three sets according to their plant height, with each set having incomplete blocks. Each incomplete block of 20 experimental lines was augmented with the random placement of two height-specific check lines (We05407 and W5579, W5579 and Ia5125, or Ia5125 and IL125b). In both field locations, experimental units were one-row plots of different lengths. Plots were 3.05 m long in NY and 3.50 m long in WI. Both locations had an inter-row spacing of 0.76 m, with a 0.91 m alley at the end of each plot. In NY, 25 kernels were planted per plot and thinned to 12 plants per plot. In WI, 12 kernels were planted in each plot, but plots were not thinned.

In all environments, multiple plants per plot were selfed pollinated, with two selfed ears collected by hand from each harvestable plot at 400 growing degree days after pollination (i.e., immature kernel stage at approximately 21 days after pollination) as earlier described (Baseggio *et al*. 2019). Immediately upon fresh harvest, the entirety of each dehusked ear was directly frozen in liquid nitrogen, followed by hand shelling of frozen kernels. To generate a representative composite kernel sample for each harvested plot, frozen kernels were equally sampled at random from both ears, bulked, and stored in 15 mL Falcon tubes at −80°C until lyophilization. A combined set of 1,524 plot samples from across all environments, with each sample consisting of three lyophilized kernels, was shipped to the Donald Danforth Plant Science Center (St. Louis, MO) for elemental analysis.

### Phenotypic Data Analysis

For each plot sample, the determination of elemental concentration by an inductively coupled plasma mass spectrometer (ICP-MS) was conducted separately for each of the three lyophilized kernels as previously described in Baxter *et al*. (2014). In short, each individual unground kernel was robotically weighed, digested in concentrated nitric acid, and measured for concentrations of aluminum, arsenic, boron, cadmium, calcium, cobalt, copper, iron, magnesium, manganese, molybdenum, nickel, phosphorus, potassium, rubidium, selenium, sodium, strontium, sulfur, and zinc with a PerkinElmer NexION 350D ICP-MS. Of these 20 elements, aluminum, arsenic, cobalt, selenium, and sodium were not further considered because their measured concentrations were at trace levels, vulnerable to contamination in the course of sample processing, and/or sensitive to interference from other sample matrix constituents (Ziegler *et al*. 2013). To limit the influence of extreme analytical outliers that could negatively affect the accurate estimation of variance components when initially fitting a mixed linear model to the raw data, the method of Davies and Gather (1993) was implemented similarly to its use in Baxter *et al*. (2014) to remove raw concentration values with greater than a conservative threshold of 15 median absolute deviations from the median concentration for a given element within each environment. If less than 1% of the values for a given element were negative, these negative values were set to missing.

The preliminarily processed raw ICP-MS dataset was more robustly screened for significant outliers by fitting a mixed linear model that allowed for genetic effects to be separately estimated from field design effects, following the procedure described in Wolfinger *et al*. (1997). The fitted mixed linear model was similar to that used by Baseggio *et al*. (2019) for the same experimental field design, with the notable exception that the model used in this study included a term to estimate within-plot kernel sample variance. This allowed for the removal of individual outlier measurements. For each elemental phenotype, the full model was fitted in ASReml-R version 3.0 (Gilmour *et al*. 2009) across locations (all four environments) or for each location separately (two environments, NY; or two environments, WI) as follows:

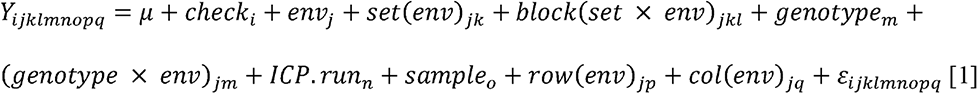

in which *Y_ijklmnopq_* is an individual phenotypic observation, μ is the grand mean, *check_i_* is the fixed effect for the *i*th check, *env_j_* is the effect of the *j*th environment, *set(env)_jk_* is the effect of the *k*th set within the *j*th environment, block*(set × env)_jkl_* is the effect of the *l*th incomplete block within the *k*th set within the *j*th environment, *genotype_m_* is the effect of the *m*th experimental genotype (non-check line), (*genotype × env)_jm_* is the effect of the interaction between the *m*th genotype and *j*th environment, *ICP.run_n_* is the laboratory effect of the *n*th ICP run, *sample_o_* is the *o*th kernel sample, *row(env)_jp_* is the effect of the *p*th plot grid row within the *j*th environment, *col(env)_jq_* is the effect of the *q*th plot grid column within the *j*th environment, and ε*_ijklmnopq_* is the heterogeneous residual error effect within each environment with a first order autoregressive correlation structure among plot residuals in the row and column directions. With the exception of the grand mean and check term, all terms were modeled as random effects. The Kenward-Roger approximation (Kenward and Roger 1997) was used to calculate degrees of freedom. Studentized deleted residuals (Neter *et al*. 1996) obtained from these mixed linear models were used to detect significant outliers for each phenotype after a Bonferroni correction (α = 0.05).

To generate best linear unbiased predictor (BLUP) values for each elemental phenotype, an iterative mixed linear model fitting procedure was conducted on the outlier-screened phenotypic dataset in ASReml-R version 3.0 (Gilmour *et al*. 2009) with the full model across locations or for each location separately. Model terms fitted as random effects including the autoregressive correlations were tested with likelihood ratio tests (Littell *et al*. 2006), followed by the removal of terms from the model that were not significant at α = 0.05. The significance of main random effects and variance component estimates are reported in Table S1. Additionally, the first order autoregressive correlation structure was statistically significant for all phenotypes. For each elemental phenotype, the final, best fitted model was used to generate a BLUP for each inbred line. The generated BLUP values were filtered to remove non-sweet corn lines, as well as sweet corn lines with the infrequent *aeduwx* or *bt2* endosperm mutations and those without available SNP marker data. This resulted in 401 sweet corn lines with more prevalent endosperm mutations [*su1*, *su1se1* (classified as *su1* for this study due to lack of informative marker genotypes), *sh2*, and *su1sh2*] that had BLUP values for elemental phenotypes across and within locations.

With variance component estimates from each best fitted model, heritability on a line-mean basis was calculated for each elemental phenotype across locations and separately for each location as previously described (Lynch and Walsh 1998; Holland *et al*. 2003; Hung *et al*. 2012). Pearson’s correlation coefficient (*r*) was used to assess the degree of association between the BLUP values of paired phenotypes. Pairwise correlations were calculated, and their significance tested at α = 0.05 with the method ‘pearson’ from the function ‘cor.test’ in R version 3.6.1 (R Core Team 2019).

### SNP Marker Genotyping

The sweet corn inbred association panel was sequenced via the genotyping-by-sequencing (GBS) procedure of Elshire *et al*. (2011) with *Ape*KI at the Cornell Biotechnology Resource Center (Cornell University, Ithaca, NY, USA) as previously described (Baseggio *et al*. 2019). The procedure of Baseggio *et al*. (2020) for SNP calling, filtering, and imputing missing genotypes was used to construct a SNP marker dataset for the genetic dissection and prediction of fresh kernel elemental phenotypes. In brief, the raw GBS sequencing data from Baseggio *et al*. (2019) was processed through the production pipeline in TASSEL 5 GBSv1 with the ZeaGBSv2.7 Production TagsOnPhysicalMap file to call SNPs at 955,690 loci in B73 RefGen_v2 coordinates (Glaubitz *et al*. 2014). These raw SNP genotype calls were merged with those of 19 sweet corn inbred lines from Romay et al. (2013) that were not included in the Baseggio *et al*. (2019) GBS dataset, allowing for the assemblage of raw SNP calls for all 401 sweet corn inbred lines with BLUP values.

The combined raw dataset was initially filtered by keeping only biallelic SNPs with a call rate > 10% and eliminating singleton (heterozygous site) and doubleton (homozygous site) SNPs that score a minor allele in only a single individual. Given the potential to have resulted from paralogous alignments, we set heterozygous genotype calls with an allele balance score (lowest allele read depth/total read depth) < 0.3 to missing. If multiple GBS samples existed for an inbred line, SNP genotype calls from samples with the same accession number/identifier were merged and discordant genotype calls set to missing if identical-by-state (IBS) values for all within-line sample comparisons were > 0.99, following the conservative IBS threshold set by Romay *et al*. (2013). A single GBS sample with the highest SNP call rate was chosen to represent an inbred line if all pairwise IBS values were less than 0.99.

The FILLIN haplotype-based imputation strategy of Swarts *et al*. (2014) was used to impute missing SNP genotypes to near completeness based on a set of maize haplotype donors with a window size of 4 kb. To improve the quality of the imputed dataset, we filtered SNPs in TASSEL 5 version 20190321 to remove those with a call rate < 70% (residual missing genotype data are expected for the haplotype-based imputation method of FILLIN; Swarts *et al*., 2014), a minor allele frequency < 5%, heterozygosity > 10%, coefficient of panmixia< 80%, or a mean read depth > 15. To uplift the genome coordinates of retained SNPs to B73 v4, the Vmatch software (Kurtz 2003) was used to align the 101 bp context sequence of each SNP to the B73 RefGen_v4 reference genome, resulting in 147,762 high-quality SNP markers scored on the 401 sweet corn inbred lines.

### Genome-Wide Association Study

A GWAS was conducted across and within locations to identify SNP markers significantly associated with each elemental phenotype following the methods of Baseggio *et al*. (2020) with minor modifications. In short, the Box-Cox power transformation (Box and Cox 1964) was used with an intercept-only model to choose the optimal value of convenient lambda (−2 to +2, 0.5 increments) (Table S2) for transforming the BLUP values of each elemental phenotype to lessen heteroscedasticity and non-normality of the residuals with the MASS package in R version 3.6.1 (R Core Team 2019). For each elemental phenotype, a mixed linear model (Yu *et al*. 2006; Zhang *et al*. 2010) that accounted for population structure and unequal relatedness with principal components (PCs) and a genomic relationship matrix (GRM; kinship) was used to test for an association between each of the 147,762 SNPs and transformed BLUP values in GEMMA software version 0.97 (Zhou and Stephens 2014). In the R package GAPIT version 2017.08.18 (Lipka *et al*. 2012), 10,773 unimputed genome-wide SNPs (call rate >90%, MAF >5%, heterozygosity <10%, coefficient of panmixia >80%, and mean read depth <15) subsampled from the complete marker dataset were used to calculate PCs with the prcomp function and the kinship matrix with VanRaden’s method 1 (VanRaden 2008). The conservative imputation of residual missing SNP genotypes as heterozygous in both marker datasets was conducted in GAPIT.

The Bayesian information criterion (BIC) (Schwarz 1978) was used to ascertain the optimal number of PCs to incorporate in the mixed linear model. Given that the predominant accumulation of some elements in the endosperm (Lombi *et al*. 2009, 2011; Pongrac *et al*. 2013; Baxter *et al*. 2014; Cheah *et al*. 2019) could potentially lead to spurious associations with *su1* and *sh2* as shown for tocotrienols and certain carotenoids (Baseggio *et al*. 2019, 2020), endosperm mutation type (*su1*, *sh2*, or *su1sh2*) was also tested with the BIC for inclusion as a covariate in the model. Of the 401 inbred lines, 384 lines had endosperm mutation type scores available from Baseggio *et al*. (2019), whereas endosperm mutation type for each of the remaining 17 lines without visual scores was predicted with the identical optimal marker-based classification models and 1000 kb marker datasets for the *su1* and *sh2* loci from Baseggio *et al*. (2019).

To approximate the amount of phenotypic variation explained by a significantly associated SNP, we calculated the difference between the likelihood-ratio-based R^2^ statistic (R^2^) of Sun *et al*. (2010) from a mixed linear model with or without the given SNP, following Baseggio *et al*. (2020). The false-discovery rate (FDR) was controlled at 5% by adjusting the *P*-values (Wald test) of SNPs tested in GEMMA using the Benjamini–Hochberg multiple test correction (Benjamini and Hochberg 1995) with the ‘p.adjust’ function in R version 3.6.1 (R Core Team 2019). Given the large variance in the estimated distance to which median genome-wide linkage disequilibrium (LD) decays to background levels (*r^2^* < 0.1 by ∼12 kb) in this association panel (Baseggio *et al*. 2019) and to account for the possibility of distant cis-regulatory elements (Ricci *et al*. 2019), candidate gene searches were limited to ± 250 kb (median *r^2^* ≤ 0.05) of the physical position of SNP markers significantly associated with an elemental phenotype. For each most plausible candidate gene, we used BLASTP to identify the top three unique best hits (E-values < 1) in Arabidopsis (Columbia-0 ecotype) and rice (*Oryza sativa* L. ssp. Japonica cv. ‘Nipponbare’) using default parameters at the TAIR (https://www.arabidopsis.org) and RAP-DB (https://rapdb.dna.affrc.go.jp) databases, respectively. The across-location (All Locs: New York, Florida, North Carolina, and Puerto Rico) results from JL analysis and GWAS of grain elemental phenotypes in the U.S. NAM panel (Ziegler *et al*. 2017) were integrated with the physical (bp) positions of GWAS signals from our study in B73 RefGen_AGPv4 coordinates following the approach of Wu *et al*. (2021).

The multi-locus mixed-model (MLMM) approach of Segura *et al*. (2012) that sequentially adds significant markers as covariates in the model was used to better clarify significant association signals with underlying large-effect loci at the level of an individual chromosome as previously described (Lipka *et al*. 2013). The optimal model was selected with the extended BIC (Chen and Chen 2008). To further assess the extent of statistical control for large-effect loci, GWAS was reconducted by including MLMM-selected SNPs as fixed effects (covariates) in mixed linear models fitted in GEMMA.

### Linkage disequilibrium

The local patterns of LD surrounding significantly associated loci were investigated by estimating pairwise LD between SNPs with the squared allele-frequency correlation (*r^2^*) method of Hill and Weir (1988) in TASSEL 5 version 20190627 (Bradbury *et al*. 2007). The marker dataset used for estimation of LD consisted of the 147,762 SNPs without imputation of the post-FILLIN residual missing SNP genotypes to heterozygotes.

### Whole-genome prediction

A univariate genomic best linear unbiased prediction (GBLUP) model (Bradbury *et al*. 2007; VanRaden 2008) was used to evaluate whole-genome prediction (WGP) on the transformed across-location BLUP values of the 15 elemental phenotypes as previously described by Baseggio *et al*. (2020). In short, the 401 line x 147,762 SNP genotype matrix with post-FILLIN missing data imputed as a heterozygous genotype was used to construct a GRM with method 1 from VanRaden (2008) in GAPIT version 2017.08.18 (Lipka *et al*. 2012). Next, the constructed GRM was modeled as a random effect to predict each individual elemental phenotype with the function ‘emmreml’ in version 3.1 of the R package EMMREML (Akdemir and Okeke 2015). Through the implementation of a five-fold cross-validation scheme conducted 50 times for each elemental phenotype, the predictive ability of a phenotype was calculated as the mean Pearson’s correlation between transformed BLUP (observed) and genomic estimated breeding values (predicted). Each fold was representative of genotype frequencies for endosperm mutants (*su1*, *sh2*, and *su1sh2*) observed in the whole association population. Endosperm mutation type (*su1*, *sh2*, or *su1sh2*) was also evaluated as a covariate in prediction models, with the same cross-validation folds used across models with or without the covariate for endosperm mutation type.

### Data availability

All raw GBS sequencing data are available from the National Center of Biotechnology Information Sequence Read Archive under accession number SRP154923 and in BioProject under accession PRJNA482446. The ZeaGBSv2.7 Production TagsOnPhysicalMap file (AllZeaGBSv2.7_ProdTOPM_20130605.topm.h5) for calling SNPs, the raw SNP genotype data in B73 AGPv2 coordinates (ZeaGBSv27_publicSamples_rawGenos_AGPv2-150114.h5) for the 19 sweet corn lines of Romay *et al*. (2013), and the maize haplotype donor file (AllZeaGBSv2.7impV5_AnonDonors4k.tar.gz) for imputing missing genotypes are on CyVerse (https://datacommons.cyverse.org/browse/iplant/home/shared/panzea/genotypes/GBS/v27). The BLUP values of the 15 elemental phenotypes and the FILLIN imputed SNP genotype calls in B73 AGPv4 coordinates for the 401 inbred lines are available at CyVerse: (https://datacommons.cyverse.org/browse/iplant/home/shared/GoreLab/dataFromPubs/Baseggio_SweetcornElement_2021). The Supplemental Figures and Tables are available at CyVerse: (https://datacommons.cyverse.org/browse/iplant/home/shared/GoreLab/dataFromPubs/Baseggio_SweetcornElement_2021/BioRxivSupplementalInformation). Except for the University of Wisconsin germplasm, all inbred lines included in the sweet corn association panel are in the public domain. A material transfer agreement is required to obtain some of the Wisconsin lines.

## RESULTS

### Phenotypic variation

The extent of phenotypic variation for 15 elements in fresh kernels as quantified by ICP-MS was evaluated in an association panel of 401 sweet corn inbred lines that was grown in two field locations (Verona, WI; and Aurora, NY) in 2014 and 2015. Of the five macroelements studied, potassium, phosphorus, sulfur, and magnesium had average concentrations greater than 1,000 μg g^−1^, whereas calcium had an average concentration of nearly 60 μg g^−1^ (Table 1). Average concentrations ranged from 0.012 (cadmium) to 24.52 (zinc) μg g for the 10 microelements. Even though cadmium had the lowest mean concentration, it had a 12.11-fold range in variation (maximum BLUP value divided by the minimum BLUP value), whereas the other 14 elements covered a 1.34- to 3.66-fold range in variation. When separating inbred lines according to their endosperm mutation type (Table 2), copper, iron, manganese, and potassium were found to be at significantly (*P* < 0.001) greater concentrations in the *sh2* (*n* = 78) group relative to the *su1* (*n* = 301) group.

**Table 1.** Means and ranges for untransformed best linear unbiased predictors (BLUPs) of 15 fresh kernel elemental phenotypes evaluated in the sweet corn association panel and estimated heritability 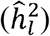 on a line-mean basis across two years and two locations.

**Table 2.** Estimated effects of endosperm mutation type from untransformed best linear unbiased predictors of 15 fresh kernel elemental phenotypes across two years and two locations.

Implying common genetic control (Baxter 2015), shared chemical and physiological properties (Marschner 2011), or storage with phytic acid (Maathuis 2009), element pairs with strong positive correlations (*r* > 0.50; *P* < 0.01) across locations (Figure S1) were as follows: strontium/calcium, magnesium/phosphorus, zinc/iron, and phosphorus/zinc. Suggestive of a distinct genetic architecture, molybdenum was the element most weakly correlated with other elements across locations (Figure S1), having a significant but very weak positive correlation (*r* = 0.12; *P* = 0.02) with zinc alone. With the exception of boron 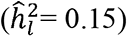 that can have elevated background levels from the use of glass (sodium borosilicate) tubes for chemical digestion (Baxter *et al*. 2014), the across-location heritability estimates (Table 1) for the elemental phenotypes were 0.40 (sulfur) and larger. The G×E interaction term was only significant for boron, copper, iron, magnesium, and rubidium, whereas the genotype term was significant for all 15 elements (Figure S2 and Table S1).

When investigating phenotypic variation between locations, we found that only the average concentration of sulfur was not significantly different between locations (Figure S3). Indicative of phenotypes with a range of responsiveness to the environment, the correlation (*r*) of elemental trait BLUPs between locations ranged from 0.08 (boron) to 0.68 (copper), with an average correlation of 0.42 (Figure S4). Despite the mostly moderate correlations between locations, within-location heritability estimates (Table S3) were comparable to those estimated across locations (Table 1) and strongly correlated (*r* = 0.81) between the NY and WI locations. Altogether, our findings suggest that there is value in exploring the genetic dissection of elemental fresh kernel phenotypes across- and within-locations.

### Genome-Wide Association Study

We investigated the genetic basis of natural variation for the concentration of 15 elements in fresh kernels from the sweet corn association panel of 401 inbred lines that had been evaluated in four environments (two years × two locations) and scored with 147,762 genome-wide SNP markers. Through an across-location GWAS conducted with a unified mixed linear model that accounted for population structure, relatedness, and endosperm mutation type, 220 unique SNPs were found to associate with one of three elements (cadmium, zinc, or boron) at a genome-wide FDR of 5% (Table S4). Significant association signals were only found on chromosomes 1 (boron), 2 (cadmium), and 7 (zinc), with the exception of a single SNP associated with cadmium on chromosome 8 (Figure S5).

The strongest association signal was identified for the concentration of cadmium, consisting of 191 significant SNPs that covered a 36.03-Mb interval within a long-range LD region of chromosome 2 (Figure 1A). The peak SNP (S2_157751802; *P*-value 1.53×10^−23^; 162,398,589 bp) for this complex association signal (Table S4), which explained 18% of the phenotypic variance for cadmium, was positioned within the open reading frame (ORF) of a gene (Zm00001d005174) that codes for a protein that belongs to the superfamily of uridine diphosphate-glycosyltransferases (www.maizegdb.org). However, this and other candidate genes within 250 kb of the peak SNP (www.maizegdb.org) were considered to unlikely be involved in cadmium accumulation. Given the extensive LD within this recombination suppressed region (Gore *et al*. 2009; Rodgers-Melnick *et al*. 2016), we searched for more plausible candidate genes within 250 kb of other significant SNPs in LD with the peak SNP. This led to our primary focus on five SNPs significantly associated with cadmium that were ∼630 kb from the peak SNP, in moderately strong LD (mean *r^2^* of 0.48) with the peak SNP and located within the *heavy metal ATPase3* (*hma3*; Zm00001d005190) and *heavy metal ATPase4* (*hma4*; Zm00001d005189) genes (Table S5). Notably, the *hma3* and *hma4* genes encode proteins with 71% and 66% amino acid sequence identity to OsHMA3 (Table S5), a P_1B_-type ATPase involved in sequestration of cadmium in root vacuoles of rice (Ueno *et al*. 2010).

**Figure 1.**
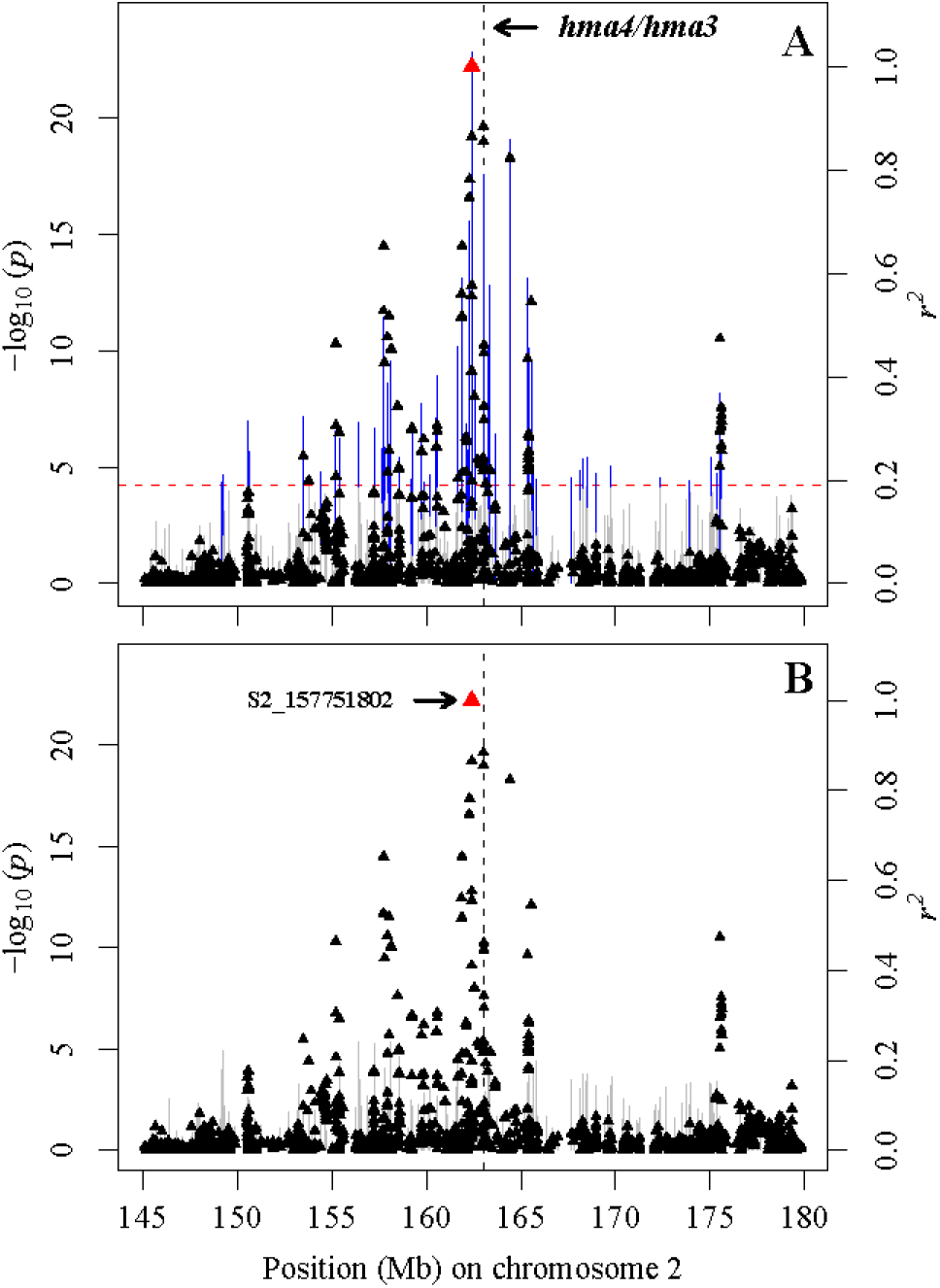
Genome-wide association study for cadmium concentration in fresh kernels of sweet corn. (A) Scatter plot of association results from a mixed model analysis and linkage disequilibrium (LD) estimates (*r^2^*). The vertical lines are –log_10_ *P*-values of single nucleotide polymorphisms (SNPs), with the blue color representing SNPs that are statistically significant at a 5% false discovery rate (FDR). Triangles are the *r^2^* values of each SNP relative to the peak SNP (indicated in red) at 162,398,589 bp (B73 RefGen_v4) on chromosome 2. The red horizontal dashed line indicates the –log_10_ *P*-value of the least statistically significant SNP at a 5% FDR. The black vertical dashed lines indicate the genomic positions of the *heavy metal ATPase4* (*hma4*; Zm00001d005189; 163,016,710-163,020,248 bp) and *heavy metal ATPase3* (*hma3*; Zm00001d005190; 163,038,225-163,041,426 bp) genes. These two genes are separated by a physical distance of ∼18 kb, thus their positions are not distinguishable at the plotted scale. (B) Scatter plot of association results from a conditional mixed linear model analysis and LD estimates (*r^2^*). The SNP from the optimal multi-locus mixed-model (S2_157751802) was included as a covariate in the mixed linear model to control for the large-effect locus. None of the tested SNPs were significant at a 5% FDR.

To better resolve the expansive association signal resulting from a large-effect locus located in a long range, high LD genomic region, a chromosome-wide multi-locus mixed-model procedure (MLMM) was conducted for cadmium. The resulting optimal model only included the peak SNP (S2_157751802) on chromosome 2 (Table S6). When GWAS was reconducted with this MLMM-selected SNP included as a covariate in the mixed linear model to control for this large-effect locus on chromosome 2, all other previously significant associations on chromosomes 2 and 8 were no longer significant at a genome-wide FDR of 5% (Figure 1B).

We identified 21 SNPs that spanned a 1.21-Mb region on chromosome 7 that were significantly associated with the concentration of zinc in fresh kernels (Figure 2A). The peak SNP (S7_174515604; *P*-value 3.19×10^−10^; 180,076,727 bp) for this association signal explained 7% of the phenotypic variance for zinc and was located within the ORF of a gene (Zm00001d022563) that encodes a tetratricopeptide repeat-like superfamily protein (www.maizegdb.org). Notably, this peak SNP was located ∼111 kb from the *nicotianamine synthase5* (*nas5*) gene (Zm00001d022557; Table S5), which encodes a class II NAS that presumably contributes to the production of the metal chelator nicotianamine (Zhou *et al*. 2013). Of the 21 detected SNP-zinc associations, SNP S7_174279369, which was located ∼160 kb from *nas5* and in moderately strong LD (*r^2^* = 0.49) with the peak SNP for zinc, also had a near significant association (FDR-adjusted *P*-value 0.06) with the concentration of iron in fresh kernels. When using the chromosome-wide MLMM procedure to better clarify the association signal complex within the 1.21-Mb region on chromosome 7 for zinc, only the peak SNP S7_174515604 was included in the optimal model (Table S6). With the MLMM-selected peak SNP as a covariate, a conditional mixed model analysis did not detect any SNPs significantly associated with zinc (Figure 2B).

**Figure 2.**
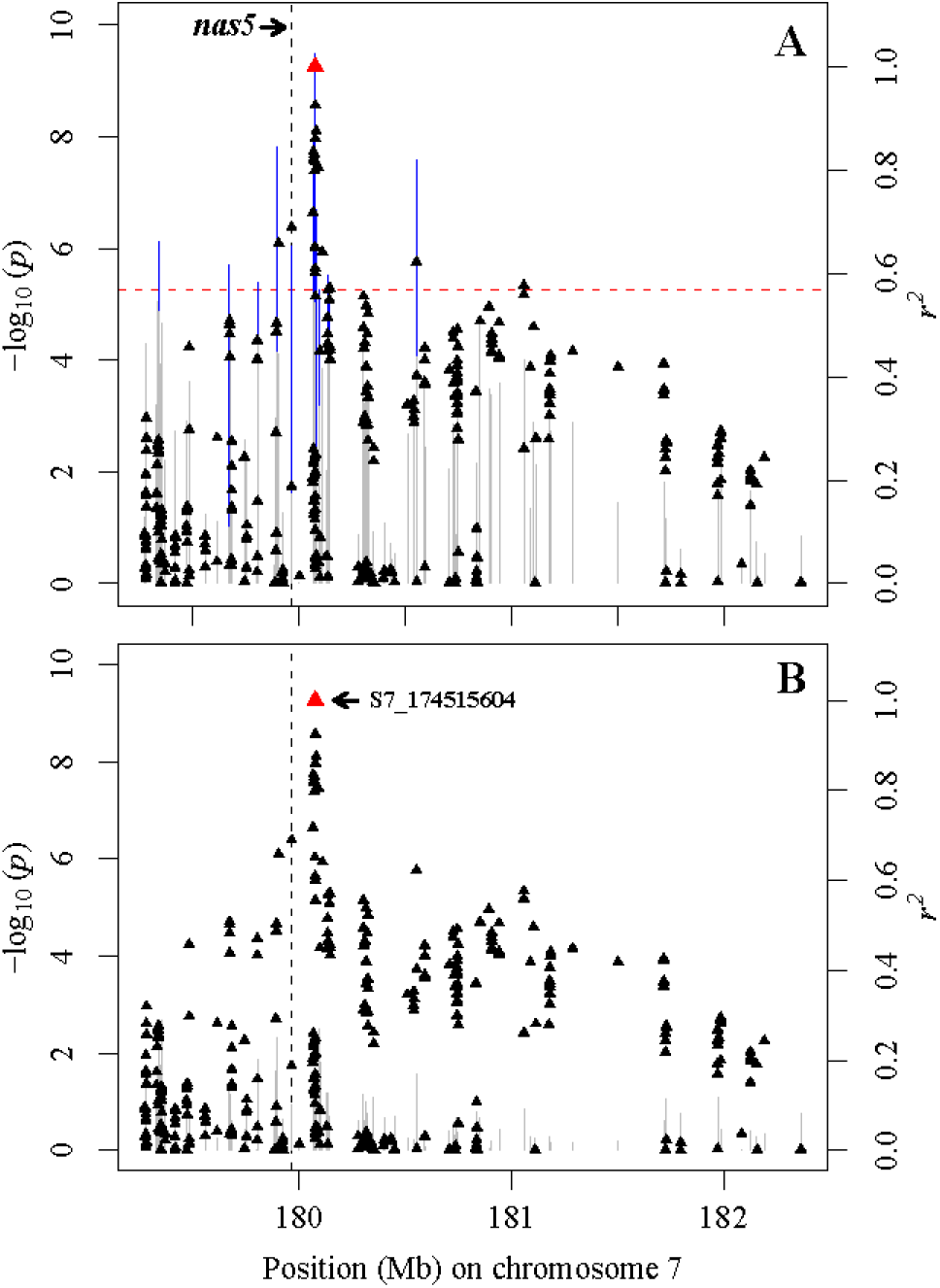
Genome-wide association study for zinc concentration in fresh kernels of sweet corn. (A) Scatter plot of association results from a mixed model analysis and linkage disequilibrium (LD) estimates (*r^2^*). The vertical lines are –log_10_ *P*-values of single nucleotide polymorphisms (SNPs), with the blue color representing SNPs that are statistically significant at a 5% false discovery rate (FDR). Triangles are the *r^2^* values of each SNP relative to the peak SNP (indicated in red) at 180,076,727 bp (B73 RefGen_v4) on chromosome 7. The red horizontal dashed line indicates the –log_10_ *P*-value of the least statistically significant SNP at a 5% FDR. The black vertical dashed line indicates the genomic position of the *nicotianamine synthase5* (*nas5*; Zm00001d022557;179,964,493-179,965,584 bp) gene. (B) Scatter plot of association results from a conditional mixed linear model analysis and LD estimates (*r^2^*). The SNP from the optimal multi-locus mixed-model (S7_174515604) was included as a covariate in the mixed linear model to control for the large-effect locus. None of the tested SNPs were significant at a 5% FDR.

Compared to cadmium and zinc, a relatively weaker association signal consisting of seven significant SNPs was identified for boron concentration on chromosome 1. Collectively, these seven SNPs comprised a 98.99-kb interval. The peak SNP (S1_189146031; *P*-value 5.01×10^−7^; 191,327,920 bp), which was located within the ORF of a gene (Zm00001d031473) encoding a protein with 87% sequence identity to an aminoacylase in rice (Table S5), explained 5% of the phenotypic variance for boron concentration. Of the other candidate genes within 250 kb of the peak SNP position, a gene (Zm00001d031476) found to be ∼45 kb away from the peak SNP encodes a protein with 40-45% amino acid sequence identity to two heavy metal-associated isoprenylated plant proteins (HIPPs) in Arabidopsis (Table S5) that are putative metallochaperones (de Abreu-Neto *et al*. 2013). Indicative of a weaker effect locus, the MLMM procedure at the chromosome-wide level did not select any SNP for the optimal model.

To identify marker-trait associations that may be location-specific, we conducted GWAS for the 15 fresh kernel elemental phenotypes within each location (NY, Figure S6; WI, Figure S7), resulting in significant association signals detected for cadmium (NY and WI), zinc (NY and WI), nickel (NY), and calcium (WI) at 5% FDR (Table S4). In the NY location, the association signal for cadmium consisted of 198 SNPs that defined a 36.89-Mb region on chromosome 2, with the peak SNP (S2_157751802; *P*-value 2.64×10^−19^; 162,398,589 bp) for the signal the same as detected for cadmium across locations (Table S4). Potentially the result of environmental variation combined with lower mapping precision from fewer evaluated inbred lines relative to the NY location (Table S3), a different peak association signal (S2_159765450; *P*-value 1.98×10^−15^; 164,415,588 bp) that contained 81 SNPs covering 32.92-Mb on chromosome 2 was identified for cadmium in the WI location (Table S4). The peak SNP for the WI location was ∼1.38-Mb from *hma3* and *hma4*, whereas the peak SNP in the NY location was ∼630 kb from the same two candidate genes. Within each location, the chromosome-level MLMM procedure was performed for cadmium (Table S6), selecting only the peak SNP that when included as a covariate in a conditional GWAS rendered all other associations on chromosomes 1 (NY), 2 (NY and WI), and 3 (NY and WI) no longer significant.

In both locations, the same SNP served as the peak association signal (NY: *P*-value 4.89×10^−8^; WI: *P*-value 7.75×10^−8^) for zinc, which was ∼111 kb from *nas5* on chromosome 7 (Table S4). The same SNP was also detected as the peak association signal for zinc across locations. The chromosome-wide MLMM approach selected only the peak SNP in the best model within each location (Table S6). With the peak SNP as a covariate in a conditional GWAS, the other previously detected SNPs for zinc on chromosome 7 in NY (6 SNPs) and WI (3 SNPs) did not remain significant. Similar to the results from conducting the across-location GWAS, SNP S7_174279369, which was ∼160 kb from *nas5*, had a weak association (FDR-adjusted *P*-value 0.07) with iron in the NY location. In contrast to the across-location GWAS, however, this SNP was not significantly associated with zinc in either the NY or WI locations (Table S4).

A significant association signal was detected for calcium in the WI location but not the NY location (Table S4). This signal for calcium consisted of four SNPs on chromosome 10. The peak SNP (S10_124069084; *P*-value 6.09×10^−08^; 125,112,781 bp) was ∼36 kb from a gene (Zm00001d025654) that codes for a protein with 47-49% sequence identity to two HIPPs in Arabidopsis (Table S5). Additionally, this peak SNP was selected by the MLMM in the optimal model at the chromosome-wide level (Table S6). In concordance with other conditional GWAS results, no other SNPs remained significant when SNP S10_124069084 was used as a covariate in the mixed linear model.

Of the two field locations, we only detected a significant association for nickel in the NY location. The association signal for nickel on chromosome 9 consisted of four SNPs, with the peak SNP (S9_2213924; *P*-value 2.80×10^−08^; 1,934,330 bp) for the signal selected by the MLMM in the optimal model (Tables S4 and S6). The peak SNP was within the ORF of a gene (Zm00001d044768) that encodes a protein 39-47% identical at the amino acid sequence level to three members of the NITRATE TRANSPORTER 1/PEPTIDE TRANSPORTER (NRT1/PTR) family (NPF) in Arabidopsis (Table S5) that transport nitrate, amino acids, and hormones (Léran *et al*. 2014). Additionally, this peak SNP was ∼44 kb from a gene (Zm00001d044771) coding a protein with 49-52% sequence identity to three matrix metalloproteinases (MMPs) in Arabidopsis (Table S5) that have a zinc-binding sequence (Marino and Funk 2012). All other significant SNPs on chromosome 9 and a single significant SNP on chromosome 5 were no longer significantly associated with nickel in a conditional mixed linear model with the peak SNP as a covariate.

### Whole-genome prediction

We evaluated the predictive ability of WGP with 147,762 SNP markers for the across-location concentrations of 15 elements that had been scored on fresh kernels of the 401 inbred lines. The 15 elements had an average predictive ability of 0.37, ranging in abilities from 0.19 for rubidium to 0.52 for copper (Table 3). The predictive abilities were above average for iron (0.45) and zinc (0.49), suggesting that genomic selection could be used to increase the concentration of both nutritionally limiting microelements in fresh sweet corn kernels. A strong positive Pearson’s correlation coefficient (*r* = 0.62 *P*-value < 0.05) was found between heritability estimates and predictive abilities for the 15 elemental phenotypes. Given the detection of significant differences among endosperm mutation types for 10 of the 15 elemental phenotypes (Table 2), endosperm mutation type was tested as an included covariate in WGP models, but changes to prediction abilities from its inclusion were zero to negligible (Table 3).

**Table 3.** Predictive abilities of whole-genome prediction models for 15 fresh kernel elemental phenotypes in the sweet corn association panel.

## DISCUSSION

Maintaining elemental homeostasis is critical for plants to realize optimal growth and complete their life cycle (Marschner 2011). Additionally, the elemental content and composition of edible plant parts are influenced by genetic and environmental factors (Watanabe *et al*. 2007; Baxter and Dilkes 2012; Baxter *et al*. 2014). Several genes responsible for natural variation of elemental levels in root and shoot tissues have been identified and characterized in plants (Huang and Salt 2016; Yang *et al*. 2018), but considerable effort remains to pinpoint the genes regulating elemental levels in seed of crops. To further this research, we examined the extent of phenotypic variation for elemental concentrations in fresh kernels and performed a GWAS to identify candidate genes controlling this phenotypic variability in a sweet corn association panel. We also evaluated the ability of genome-wide markers to predict elemental concentrations, providing insights into the potential of genomic selection for optimizing the elemental profile of fresh kernels for human health and nutrition, especially iron and zinc. To the best of our knowledge, this work is the most extensive quantitative genetic analysis of elemental concentrations in fresh kernels of sweet corn.

The rank order of average concentrations for the measured elements in fresh kernels from the sweet corn association panel was highly concordant with that observed for the same elements in physiologically mature grain from the maize Ames panel of non-sweet corn tropical and temperate inbred lines (Wu *et al*. 2021) and the B73 (dent; *Su1*) x IL14H (sweet corn; *su1*) recombinant inbred line (RIL) family of the U.S. NAM panel (Baxter *et al*. 2014). Our sweet corn association panel showed a range of 13.66-28.11 and 18.58-32.10 μg g^−1^ on a dry weight basis for iron and zinc, respectively. However, these ranges were both lower and narrower than that found for iron (14.62 - 36.33 μg g^−1^; 3.29 S.D.) and zinc (12.59 - 52.32 μg g^−1^; 4.36 S.D.) in physiologically mature grain from the maize Ames panel (Wu *et al*. 2021). With consideration of losses from processing and bioavailability, the recommended target level iron and zinc grain concentrations established by HarvestPlus, which has a primary focus on developing nations where nutritional deficiencies are prevalent, are 60 and 38 μ g dry weight, respectively, for developing biofortified maize based on achieving ∼30-40% of estimated average requirements for adult women (nonpregnant, nonlactating) when consuming 400 g d^−1^ of whole maize grain (Bouis and Welch 2010). Although the maximum iron concentration observed in our sweet corn association panel (28.11 μg g^−1^ dry weight) is ∼2-fold lower than the HarvestPlus breeding target, there are six inbred lines that have zinc concentrations ranging from 30.00 to 32.10 μg g^−1^ dry weight. As it relates to the consumption of fresh sweet corn when not a primary source of daily calories, the estimated maximum fresh kernel concentrations observed in our association panel would provide approximately 4-9% and 7-10% of the RDA of iron and zinc, respectively, for adult non-elderly women (nonpregnant, nonlactating) and men when consuming 100 g (∼75% water) of uncooked fresh sweet corn. Irrespective of lacking experimental data from bioavailability assays, our comparison of phenotypic distributions to dietary guidelines implies that the top 5% highest ranking lines for iron (≥ 22 μg g^−1^ dry weight) and zinc (≥ 28 μg g^−1^ dry weight) concentrations have promise for establishing a biofortification program for sweet corn.

We assessed whether the concentrations of elements differed significantly among endosperm mutation group types. Of the 15 elements, copper, iron, manganese, potassium, and sulfur were highly significantly different (*P* < 0.001) between two or more endosperm mutation type groups (*su1*, *sh2*, and *su1sh2*) (Table 2). Relatedly, Baxter *et al*. (2014) showed that the content for four (iron, manganese, potassium, and sulfur) of these five elements significantly differed (*P* < 0.0005) between visibly ‘wrinkled’ (*su1*/*su1*) and ‘non-wrinkled’ (*Su1*/*su1* or *Su1*/*Su1*) kernels harvested at physiological maturity. As it relates to the spatial distribution of elements in sweet corn kernels, Cheah *et al*. (2019) analyzed immature (21 DAP) kernels from a single sweet corn (*sh2*) variety via synchrotron-based X-ray fluorescence microscopy to reveal that potassium and calcium were generally present throughout the kernel, sulfur concentrated mainly in the axis of the embryo and the periphery of the endosperm, and the scutellum of the embryo had at least 20-fold higher concentrations of phosphorus, iron, zinc, and manganese than in the endosperm. Notably, Cheah *et al*. (2019) also showed that these spatial distribution maps for elements were highly similar to those of immature maize (non-sweet corn) kernels. Despite these valuable insights from earlier studies, further experimental work will be needed to determine whether the observed significant difference in concentrations of the five elements among endosperm mutation type groups in our sweet corn association panel were attributed to variation in physiological, genetic, and/or physical attributes of fresh kernels.

Conducting GWAS across locations for the concentrations of elements in fresh kernels of the sweet corn association panel resulted in the identification of candidate genes associated with cadmium and zinc at the genome-wide level. Of these elements, the strongest association signal was for cadmium, having an association signal on chromosome 2 that spread more than 35-Mb across a recombinationally inert genomic region. Additionally, the peak SNP of this association signal co-localized with the single across-location QTL detected for grain cadmium concentration in the maize NAM panel (Tables S4 and S7), but the GWAS resolution for this region in the NAM panel (Table S8) was too limiting to convincingly identify an underlying causal gene (Ziegler *et al*. 2017). In our sweet corn association panel, however, two likely candidate causal genes were identified, *hma3* and *hma4*, both having SNPs in moderately strong LD with the peak SNP of the association signal. These genes are two of a 12-member gene family encoding HMAs in the genome of maize inbred line B73 (Cao *et al*. 2019).

Of the 12 HMA genes, the proteins encoded by *hma3* and *hma4* have high sequence identity (Table S5) to the P_1B_-type ATPase, OsHMA3—a tonoplast-localized zinc/cadmium transporter that has been shown to be expressed in rice roots, mediates cadmium and zinc vacuolar sequestration and, as such, participates in zinc homeostasis and root-to-shoot cadmium translocation (Ueno *et al*. 2010; Miyadate *et al*. 2011; Sasaki *et al*. 2014; Cai *et al*. 2019). The loss-of-function of *OsHMA3* has been associated with cadmium accumulation in rice grains, whereas low-cadmium rice cultivars express a functional *OsHMA3* (Ueno *et al*. 2010). Furthermore, Ueno *et al*. (2010) have shown that the overexpression of *OsHMA3* selectively decreased the accumulation of cadmium, but not other elements in the grain. In a follow-up study, Sasaki *et al*. (2014) showed that overexpression of *OsHMA3* resulted in sequestration of both cadmium and zinc in rice root vacuoles, but the concentration of zinc in shoots was unaffected through the constitutive upregulation of transporter genes having putative involvement in the uptake and translocation of zinc. In agreement with Sasaki *et al*. (2014), we found no evidence of the large-effect locus spanning *hma3* and *hma4* having a significant association with zinc concentration in fresh sweet corn kernels. Importantly, Cao *et al*. (2019) identified several polymorphisms within *hma3* to be significantly associated with leaf cadmium concentration at the seedling and adult plant stages in a maize diversity panel. Moreover, they further showed the expression level of *hma3* to be highly upregulated in the roots of B73 in response to cadmium stress, whereas the expression of *hma4* was undetectable in roots under the same conditions (Cao *et al*. 2019). Considering this, we propose that *hma3* is the more likely of the two genes to have played a key genetic role in the accumulation of cadmium in fresh kernels.

The *nas5* gene was found to be within 250 kb of the peak SNP for the across-location zinc and iron association signals on chromosome 7. These findings co-locate with the GWAS results of Ziegler *et al*. (2017) and Wu *et al*. (2021) from the U.S. maize NAM (Tables S4, S7, and S8) and Ames mapping panels, respectively, that implicated *nas5* as a possible pleiotropic controller for the concentrations of both zinc and iron in physiologically mature grain samples. As one of nine gene family members in the B73 reference genome, *nas5* phylogenetically groups together with *nas3* and *nas4*, which together comprise class II *nas* genes (Zhou *et al*. 2013). The NAS enzyme encoded by *nas5* is hypothesized to be involved in the production of the non-proteinogenic amino acid, nicotianamine, an efficient chelator of transition metals including zinc and iron (Takahashi *et al*. 2003; Curie *et al*. 2009; Swamy *et al*. 2016). In addition to a suggested role in intracellular metal homeostasis, nicotianamine facilitates phloem-based metal delivery from source (e.g., leaves) to sink (e.g., seeds) tissues (Takahashi *et al*. 2003; Curie *et al*. 2009; Swamy *et al*. 2016). Nicotianamine also serves as a precursor for the synthesis of root-exuded mugineic acid-type phytosiderophores that chelate divalent metals for eventual root uptake (Curie *et al*. 2009; Swamy *et al*. 2016).

Consistent with a proposed role of *nas5* in long-distance metal transport rather than uptake into roots, Zhou *et al*. (2013) showed that transcripts of class II *nas* genes including *nas5* accumulated mainly in maize leaves and sheath, whereas class I *nas* genes were predominantly expressed in maize roots. Interestingly, of the three class II *nas* genes*, nas5* was more highly expressed in maize stems, further suggesting its contribution to long-distance metal transport and perhaps its contribution to metal loading to seeds (Zhou *et al*. 2013). In addition, the transcriptional expression level of *nas5* was downregulated by iron deficiency in both shoots and roots but upregulated under excess iron and zinc in roots. Activation tagging of *OsNAS3*, the closest rice homolog of *nas5* (Zhou *et al*. 2013), produced an increased level of nicotianamine that resulted in elevated levels of zinc and iron in shoots, roots, and seeds of activation-tagged rice plants (Lee *et al*. 2009). Despite the lack of functional validation, results from the maize association mapping and transgenic rice studies strongly support the nomination of *nas5* as a causal gene for controlling zinc and iron concentrations in fresh sweet corn kernels.

Similar to our findings with *nas5* in the sweet corn association panel, Wu *et al*. (2021) also identified a stronger association signal of *nas5* with zinc relative to iron in the maize Ames panel. This finding is not entirely surprising considering that the affinity constants (Kd) of complexes of nicotianamine with zinc is higher than for nicotianamine with iron (Curie *et al*. 2009; Gayomba *et al*. 2015). Therefore, it is plausible to propose that equimolar concentrations of iron and zinc would result in the selection of zinc for nicotianamine over iron (Curie *et al*. 2009; Gayomba *et al*. 2015). It is also noteworthy that the concentration of zinc in the phloem sap is thought to be higher than the concentration of iron (Reviewed in Gayomba *et al*. 2015), thus reinforcing the suggested role of nicotianamine and *nas5* in zinc accumulation in fresh kernels.

In contrast to the highly probable causality of the *hma3* and *nas5* loci detected via GWAS across and within both locations, there is only moderately compelling evidence for the genetic involvement of identified candidate genes for the concentrations of boron (across locations), calcium (WI), and nickel (NY). Of the genes within 250 kb of the peak SNP for the boron association signal on chromosome 1, two genes encoding a putative protein with sequence identity to either an aminoacylase (Zm00001d031473) or HIPP (Zm00001d031476) were the most plausible candidates. Under boron deficiency, aminoacylases (metalloenzymes involved in amino acid metabolism) have been shown to have decreased protein and increased transcript levels in *Brassica napus* L. roots and *Citrus sinensis* leaves, respectively (Wang *et al*. 2010; Lu *et al*. 2015). Transcriptome profiling revealed a *HIPP* to be upregulated in leaves and roots of black poplar (*Populus nigra* L.) grown under boron toxicity (Yıldırım and Uylaş 2016), which perhaps is not surprising given that HIPPs are metallochaperones involved in the transport of metallic ions and response to abiotic stresses (de Abreu-Neto *et al*. 2013). Despite these findings in other plant systems, the exact mechanism by which the identified aminoacylase and HIPP would have contributed to the accumulation of boron in fresh kernels is unknown but nevertheless merits further experimental investigation.

Although the peak SNP associated with calcium concentration in the WI location resided within a genomic region that lacked a definitive candidate gene, it was located ∼36 kb from a gene (Zm00001d025654) encoding a putative HIPP. However, to our knowledge, HIPPs have never been experimentally shown to bind Ca^2+^ (de Abreu-Neto *et al*. 2013), thus an *in vitro* study would be needed to determine whether Ca^2+^ is bound by the putative HIPP that Zm00001d025654 encodes. It is interesting that collectively, two different HIPP candidate genes were identified for the concentrations of boron and calcium, but these proteins would not be expected to have similar roles given that calcium and boron have different chemical and physiological properties (Marschner 2011). Regardless, these findings open new avenues of inquiry that could deepen our understanding of the genetic basis of boron and calcium accumulation in fresh sweet corn kernels.

In contrast to calcium, the nickel association signal for the NY location coincided with association signals detected for nickel grain concentrations in the maize NAM (Tables S4, S7, and S8) and Ames panels (Ziegler *et al*. 2017; Wu *et al*. 2021). This still genetically unresolved signal consisted of two possible candidate genes, an MMP (Zm00001d044771) and NPF member (Zm00001d044768). Even though MMPs could conceivably bind nickel in place of zinc (Cerdà-Costa and Gomis-Rüth 2014), their proteolytic activities to remodel the extracellular matrix (Marino and Funk 2012) lack a clear connection to nickel transport or accumulation. Additionally, the putative NPF member encoded by Zm00001d044768 is hypothesized to transport nitrate given its high sequence identity to members of the NPF5 subfamily in Arabidopsis (Table S5) (Niño-González *et al*. 2019), thus reducing but not completely eliminating the possibility that this yet-to-be characterized protein transports nickel or a substrate that binds nickel.

We showed that, on average, moderate predictive abilities were achieved through the application of WGP for the across-location concentrations of the 15 elements in fresh sweet corn kernels. Additionally, these predictive abilities were found to be strongly correlated with heritability estimates, which coheres with expectations (Combs and Bernardo 2013). In accordance with these results, Wu *et al*. (2021) analyzed 11 of these 15 elements in mature grain samples from the maize Ames association panel that had been evaluated in a single location across two years, finding that the moderate predictive abilities of the 11 elements from WGP had a strong correlation with their heritabilities. Also, the prediction abilities presented in Table 3 suggest that endosperm mutation type did not need to be considered as a covariate in WGP models to capture the genetic differences between the three groups at the genome-wide marker density employed in this study. Given that we observed significant G × E interaction for boron, copper, iron, magnesium, and rubidium, accounting for G × E in WGP models could result in slightly improved predictive abilities for these five elements. In support of this supposition, a multi-environment model incorporating G× E resulted in higher average prediction abilities for the concentration of zinc in kernels from a tropical maize inbred panel and a double haploid population compared to those from single-environment models (Mageto *et al*. 2020). Therefore, this should be an area of further exploration when conducting multi-environment genomic selection for elemental phenotypes whether at the immature or mature stages of kernel development.

The results from GWAS can be used to better inform how best to successfully implement genomic selection for the concentrations of elements in fresh sweet corn kernels. Apart from cadmium (18%) and zinc (7%), which each had a single locus explaining more than 5% of the phenotypic variance in the across-location GWAS, we observed relatively weaker association signals for boron, iron, and the other 11 elements in fresh kernels. Furthermore, we did not identify all of the strong association signals for the grain concentrations of boron, copper, manganese, molybdenum, nickel, and zinc that had been detected in the maize Ames panel (Wu *et al*. 2021). Compared to the Ames panel study of Wu *et al*. 2021 that had more than 2,000 maize inbred lines, it is likely that the size of the sweet corn association panel in our study was too underpowered to identify these loci, whether because of their lower allele frequencies and/or weaker effects. Regardless, we posit that these elemental phenotypes are generally more polygenic than carotenoid and tocochromanol levels in fresh kernels of sweet corn that have a more oligogenic inheritance (Baseggio *et al*. 2019, 2020), thus making elemental phenotypes less tractable for genetic dissection in the sweet corn association panel. Therefore, genomic selection is more advisable than marker-assisted selection as a breeding approach for selecting for the concentration of elements in fresh kernels (Lorenz *et al*. 2011; Desta and Ortiz 2014). However, it is still worthwhile to assess the inclusion of large-effect loci as fixed effects such as those for cadmium and zinc in WGP models, as it could result in higher prediction abilities in specific sweet corn breeding populations (Bernardo 2014).

## Conclusions

We used a sweet corn association panel to study the quantitative genetics of natural variation for the concentrations of 15 elements in fresh kernels. Through an across-location GWAS, we strongly implicated the candidate causal genes *nas5* with iron/zinc and *hma3* with cadmium. Given that iron and zinc accumulation in fresh kernels have a partially shared genetic basis, the genetic correlation between these two phenotypes can be leveraged with multi-trait genomic selection approaches to possibly exceed the prediction accuracy of single-trait genomic selection (Jia and Jannink 2012) for simultaneous genetic gains in zinc and iron concentrations. Such efforts would help to address iron and zinc deficiencies of women, children, and older adults in the U.S. (Clark 2008) where sweet corn is highly consumed as a fresh vegetable. Importantly, the across- and within-location association signals at the *nas5* and *hma3* loci were specific to zinc/iron and cadmium, respectively. This suggests that genomic selection for lower cadmium accumulation to reduce possible toxicity should not influence zinc accumulation in the kernel. Even though 100 g of fresh sweet corn (medium-sized ear) with the maximum concentration of cadmium found in this panel is estimated to provide less than 2% of the provisional tolerable intake for this element in a day (0.8 μg/kg bw/day) (JECFA 2011) when consumed by a 70 kg person, efforts should be dedicated towards developing haplotype tagging SNP markers at the *nas5* and *hma3* loci for breeding sweet corn that has lower cadmium but higher bioavailable zinc and iron, considering that sweet corn can be grown in regions with naturally elevated cadmium levels, or with low zinc and iron levels.

## Author Contributions

M.B., O.K.V., and M.A.G. coLwrote the manuscript; M.B. led the data analysis; G.Z. performed the ICP-MS analyses and elemental quantifications; N.K. and M.M. provided overall management of panel growth (planting, pollination, harvesting); M.B., M.M., and J.C. generated the marker datasets; I.B. oversaw the ICP-MS analyses and provided biological interpretation; C.R.B. and J.P.H. uplifted the NAM JL-GWAS results; D.W. integrated and provided biological interpretation of the NAM JL-GWAS results; O.K.V. provided biological interpretation of the candidate gene associations; W.F.T., M.M., and E.S.B. constructed the association panel; E.S.B., M.E.S., W.F.T., and M.A.G. conceived and designed the project; M.A.G. oversaw the data analysis, project management, design, and coordination.

## Acknowledgements

This research was supported by the National Institute of Food and Agriculture (NIFA); the USDA Hatch under accession numbers 100397 (M.A.G.), 1010428 (M.A.G.), 1013637 (M.A.G.), 1013641 (M.A.G.), and 142 AAC6861 072600 4 (W.F.T.), HarvestPlus (M.A.G.), the National Science Foundation (IOS□1546657 to C.R.B. and M.A.G.), USDA-NIFA grant numbers 2021-67013-33798 (O.K.V.) and 2018-67013-27418 (O.K.V.), Cornell University startup funds (M.A.G.), and the USDA□ARS (E.S.B. and I.B.). We are grateful to Jenna Hershberger for providing valuable feedback on an earlier version of the manuscript. Mention of trade names or commercial products in this publication is solely for the purpose of providing specific information and does not imply recommendation or endorsement by the USDA. The USDA is an equal opportunity provider and employer. We thank the current and past members of the Tracy and Gore labs for their efforts in pollination, harvest, and sample preparation. We also thank Lily Hislop and Patrick Flannery for sharing information from the Wisconsin field trials.

## Table Captions

**Table S1.** Sources of variation for 15 elemental phenotypes in fresh sweet corn kernels from across locations, only NY, and only WI. The bolded variance component estimates are for significant random effect terms according to a likelihood ratio test (α = 0.05).

**Table S2.** Lambda values used in Box-Cox transformation of 15 fresh kernel elemental phenotypes in sweet corn.

**Table S3.** Means and ranges for best linear unbiased predictors (BLUPs) of 15 fresh kernel elemental phenotypes evaluated in the sweet corn association panel and estimated heritability on a line-mean basis across two years in each of two locations.

**Table S4.** Significant results from a genome-wide association study of 15 fresh kernel elemental phenotypes in sweet corn and their intersection with joint-linkage QTL support intervals (NAM QTL number; Table S7) of elemental phenotypes analyzed in the maize NAM panel (Ziegler *et al*. 2017).

**Table S5.** List of top BLASTP hits in rice and Arabidopsis for the eight maize candidate genes identified via a genome-wide association study of 15 fresh kernel elemental phenotypes in sweet corn.

**Table S6.** Multi-locus mixed-model results from an analysis of fresh kernel elemental phenotypes for chromosomes 2, 7, 9, and 10.

**Table S7.** Joint-linkage QTL support intervals (SI) of elemental phenotypes analyzed in the maize NAM panel (Ziegler *et al*. 2017) uplifted from B73 RefGen_v2 to v4.

**Table S8.** Genome-wide association study results of elemental phenotypes analyzed in the maize NAM panel (Ziegler *et al*. 2017) uplifted from RefGen_v2 to v4. Only NAM marker variants with resample model inclusion probability (RMIP) ≥ 5 are shown and those that reside within joint-linkage QTL support intervals (Table S7) are demarcated in the “NAM QTL number” column. The relationship of NAM marker variants to the four candidate genes identified in genome-wide association study in sweet corn that are coincident with joint-linkage QTL support intervals are presented.

## Figure Legends

**Figure S1.** Correlation matrix for untransformed, across-location BLUPs of 15 elemental phenotypes from fresh kernels in the sweet corn association panel. Pearson’s correlation coefficients (*r*) are presented in the upper triangle, while the corresponding *P*-values for the significance of associations (α = 0.05) are displayed below the diagonal.

**Figure S2.** Sources of variation for 15 elemental phenotypes in fresh sweet corn kernels from across locations (A), only NY (B), and only WI (C). The phenotypic variance of each trait was statistically separated into the following components: environment (Env), set within environment [Set(Env)], block within set within environment [Block(Set×Env)], genotype (Geno), genotype-by-environment interaction (Geno×Env), inductively coupled plasma mass spectrometry run (ICP), kernel sample (Sample), row within environment [Row(Env)], column within environment [Col(Env)], and residual error variance (Residual). Variance component estimates were calculated for all random effects from the full model (Equation 1).

**Figure S3.** Distribution of within-location BLUP values for 15 fresh kernel elemental phenotypes in the sweet corn association panel evaluated in NY and WI. Mean values of fresh kernel elemental concentration with significant differences (*P* < 0.05) between the two locations were designated with an asterisk (‘*’) according to a paired *t*-test.

**Figure S4.** Correlation matrix for untransformed, within-location BLUPs of 15 elemental phenotypes from fresh kernels in the sweet corn association panel. Pearson’s correlation coefficients (*r*) between phenotypes within NY and WI are presented in the upper and lower triangles, respectively. The diagonal represents the correlation of each trait between locations. Bolded values correspond to significant correlations (*P*-value ≤ 0.05).

**Figure S5.** Genome-wide association study conducted across both locations for 15 fresh kernel elemental phenotypes in sweet corn. Each point represents a SNP with its −log_10_ *P*-value (y-axis) from a mixed linear model analysis plotted as a function of physical position (B73 RefGen_v4) across the 10 chromosomes of maize (x-axis). The red horizontal dashed line indicates the −log_10_ *P*-value of the least statistically significant SNP at 5% false discovery rate.

**Figure S6.** Genome-wide association study conducted within the New York location for 15 fresh kernel elemental phenotypes in sweet corn. Each point represents a SNP with its −log_10_ *P*-value (y-axis) from a mixed linear model analysis plotted as a function of physical position (B73 RefGen_v4) across the 10 chromosomes of maize (x-axis). The red horizontal dashed line indicates the −log_10_ *P*-value of the least statistically significant SNP at 5% false discovery rate.

**Figure S7.** Genome-wide association study conducted within the Wisconsin location for 15 fresh kernel elemental phenotypes in sweet corn. Each point represents a SNP with its −log_10_ *P*-value (y-axis) from a mixed linear model analysis plotted as a function of physical position (B73 RefGen_v4) across the 10 chromosomes of maize (x-axis). The red horizontal dashed line indicates the −log_10_ *P*-value of the least statistically significant SNP at 5% false discovery rate.

